# Proxies of CRISPR-Cas9 activity to aid in the identification of mutagenized Arabidopsis plants

**DOI:** 10.1101/637314

**Authors:** Renyu Li, Charles Vavrik, Cristian H. Danna

## Abstract

CRISPR-Cas9 has become the preferred gene editing technology to obtain loss-of-function mutants in plants, and hence a valuable tool to study gene function. This is mainly due to the easy reprograming of Cas9 specificity using customizable small non-coding RNAs, and to the ability to target several independent genes simultaneously. Despite these advances, the identification of CRISPR-edited plants remains time and resource consuming. Here, based on the premise that one editing event in one locus is a good predictor of editing event/s in other locus/loci, we developed a CRISPR co-editing selection strategy that greatly facilitates the identification of CRISPR-mutagenized Arabidopsis plants. This strategy is based on targeting the gene/s of interest simultaneously with a proxy of CRISPR-Cas9-directed mutagenesis. The proxy is an endogenous gene whose loss-of-function mutation produces an easy-to-detect visible phenotype that is unrelated to the expected phenotype of the gene/s under study. We tested this strategy via assessing the frequency of co-editing of three functionally unrelated proxies. We found all three proxies predicted the occurrence of mutations in either or both of the other two proxies with efficiencies ranging from 40% to 100%, dramatically reducing the number of plants that need to be screened to identify CRISPR mutants. This selection strategy provides a framework to facilitate gene function studies of gene families as well as the function of single copy genes in polyploid plant species where the identification of multiplex mutants remains challenging.

## Introduction

The ability to precisely edit genes is revolutionizing our ability to study gene function. Due to its genetic tractability, assembled genome sequence and collections of mutants, the model plant *Arabidopsis thaliana* is the preferred plant species to uncover gene function. Arabidopsis is a diploid species with a genome size of ∼135 Mb and ∼29,000 genes that encode for ∼37,000 proteins. However, in spite of the three decades of effort by many research groups, less than 40% of the Arabidopsis genes have been experimentally linked to at least one phenotype (Li *et al.* 2012; Akiyama *et al.* 2014; Provart *et al.* 2016). Forward genetic screens of chemically (i.e. EMS) or physically (i.e. γ irradiation) mutagenized Arabidopsis have been very successful at linking genes to protein function and phenotypes (Koornneef and Meinke 2010). However, reverse genetic screens of T-DNA insertional lines or transposon mutagenized lines have shown more limited success, as most single gene mutants do not produce observable phenotypes (Lloyd and Meinke 2012). Although a diploid species, the Arabidopsis genome contains a large number of gene duplications (The Arabidopsis Genome Initiative *et al.* 2000) that often translate into completely or partially redundant gene functions (Vision *et al.* 2000; Bowers *et al.* 2003). Therefore, several months of crossing single-gene mutants and selecting for multiplex is often needed to define the function of the gene/s under study.

Three gene targeting methods, Zinc Finger Nuclease (ZFN), Transcription Activator-Like Effector Nuclease (TALEN), and CRISPR-Cas9 have accelerated gene function discovery in the past few years. All three methods have been successfully used in Arabidopsis, and other plants, to generate loss-of-function mutations in specific genes (Miller *et al.* 2007, 2011; Christian *et al.* 2010; Li *et al.* 2013). All three methods are based on directing the activity of a DNA endonuclease to a specific target sequence in the genome to generate a double strand break (DSB). This is followed by the repair of the DSB via Non-Homologous End-Joining (NHEJ) (Puchta 2005; Schiml and Puchta 2016). This error-prone repair pathway generates indels that shift the frame of the coding sequence, which will likely eliminate the function of the protein encoded. While ZFN and TALEN DNA-binding specificity is based on DNA-binding protein domains that need to be optimized for every gene of interest, the gene specificity of CRISPR-Cas9 is provided by an easily customizable small guide RNA (sgRNA). Each sgRNA has three important elements: 1) a 20 bp gene-specific CRISPR RNA (crRNA) that defines the target sequence; 2) a Protospacer Adjacent Motif (PAM) site, which consists of the sequence 5’-NGG-3’ for the most commonly used *Streptococcus pyogenes* Cas9 enzyme; 3) a constant trans-activating crRNA (tracrRNA) that mediates the formation of the sgRNA-Cas9 complex (Jinek *et al.* 2012; Karvelis *et al.* 2013). Hence, targeting a new gene with CRISPR-Cas9 only requires the identification of a 20nt gene-specific sequence 3-6 bp upstream of a PAM site, and the PCR-mediated or *in vitro* DNA synthesis of the crRNA-PAM-tracrRNA sequence.

The expression of CRISPR-Cas9 in Arabidopsis and other plant species necessitates the insertion of foreign DNA encoding Cas9 and sgRNAs into the plant’s genome. For this, both Cas9 and sgRNAs are cloned in a binary plasmid and transgenic plants are obtained via *Agrobacterium tumefaciens*-mediated transformation of immature flowers (Clough and Bent 1998). In most vectors, Cas9 expression is typically controlled by a ubiquitous constitutive promoter such as UBQ10 or CaMV-35S. Agrobacterium inserts the T-DNA randomly in the genome of a few cells, most of which are somatic cells. If some germline cells are transformed with the T-DNA, the transgene will pass to the next generation (T1 generation) (Hood et al., 1998). From the few plants in the progenies that receive the T-DNA, a yet smaller subset would harbor the T-DNA in a region of the genome that allows Cas9 expression. The Cas9-expressing T1 plants, will edit the target gene/s throughout their lifetime in somatic cells as well as in some germline cells allowing CRISPR-editing to pass to the next generation (T2 progenies) (Feng *et al.* 2014). Finally, a small proportion of these T2 progenies will be homozygous for the mutation of interest. Usually, as the gene/s under study does not produce a known phenotype, the identification of the CRISPR-mutagenized T2 plants requires the non-biased DNA genotyping of a large number of plants (Kim *et al.* 2014)(Thomas *et al.* 2014)(Peng *et al.* 2018). A few alternatives have been pursued to alleviate the cost and to reduce the time invested in the identification of CRISPR-mutagenized plants. One such strategy is the use of Cas9 fusions to fluorescent proteins (i.e. GFP, mCherry, etc.) or protein tags (i.e. HA, FLAG, etc) to identify T1 plants that express CRISPR-Cas9 (Osakabe *et al.* 2016). These strategies inform about Cas9 expression in T1 plants but do not alleviate the burden of screening for CRISPR-edited plants, and typically requires imaging T1 plants under UV light or detecting Cas9 protein fusions via western blot. Therefore, the use of proxies of CRISPR-Cas9 activity instead of Cas9 expression, could significantly reduce the time and effort invested in identifying CRISPR-mutagenized plants.

Recent studies reported that CRISPR-Cas9 can simultaneously edit multiple loci (co-editing) at high frequency in the somatic tissues of T1 Arabidopsis and rice plants (Ma *et al.* 2015; Yan *et al.* 2016; Zhang *et al.* 2016; Minkenberg *et al.* 2017). Hence, we hypothesized that we could take advantage of this high co-targeting frequency to aid in the selection of CRISPR-mutagenized plants. The rationale follows: if we targeted a gene that produces a visible and easy to detect phenotype, we could use it as a proxy to identify T2 plants where other loci of interest were simultaneously edited. To test this hypothesis, we chose three genes with independent functions and located in different chromosomes as potential proxies of CRISPR-Cas9 activity, namely *GLABRA-1* (*GL1*), *Jasmonic Acid Resistant-1* (*JAR1*) and *Ethylene Insensitive-2* (*EIN2*). In Arabidopsis, the formation of leaf trichomes is contingent to the function of *GL1*. Loss-of-function mutants of *GL1* do not produce trichomes, a phenotype that is easily observable as these plants have smooth leaves (Herman and Marks 1989; Marks and Feldmann 1989). Loss-of-function mutations in *JAR1* and *EIN2* produce insensitivity to jasmonic acid (JA) and ethylene (ET), respectively. The responses to both JA and ET can be monitored in seedlings exposed to JA or ET in tissue culture plates (Alonso *et al.* 1999; Staswick *et al.* 2002). In plants harboring a wild type allele of *JAR1*, exposure to JA causes root growth inhibition. In seedling harboring a wild type allele of EIN2, exposure to ET causes hypocotyls to bend downwards. We used these three genes to test whether they are effective proxies of CRISPR-Cas9 activity, and could therefore facilitate the isolation of CRISPR-mutagenized T2 plants. Our data show that co-editing of *GL1, EIN2 or JAR1* predicts co-editing of the gene of interest with frequencies ranging from 40-100%, greatly reducing the time and resources needed to identify CRISPR-Cas9 multiplex mutants for the genes of interest. Importantly, the selection strategy laid out in this study could accelerate the process and reduce the cost of identifying multiplex mutants in plant species with large and polyploid genomes.

## Material and Methods

### Design and synthesis of sgRNA expression cassettes

The sgRNAs were designed with an online web tool at the Zhang lab (crispr.mit.edu). The occurrence of the crRNA sequences retrieved by the software was verified via PCR amplification on genomic DNA from wild-type (Col-0) plants and Sanger sequencing of the PCR amplicon. Each candidate crRNA sequence was evaluated based on the calculated specificity score and the number of off-target sites. The crRNA target site was inserted into an in-silico cloning construct template between the AtU6P promoter sequence and the tracrRNA sequence. The AtU6 promoter, crRNA, tracrRNA, and a poly-T tail together constitute a complete sgRNA expression cassette (Peterson et al., 2016). Each of the three individual cassettes was assembled into a stackable array. A 32-nucleotides sequence upstream of the first AtU6 promoter and a 17-nucleotide sequence downstream of the last poly-T tail of the array was included to facilitate future cloning (Figure S1). The final DNA sequence was synthesized through Genscript™ Custom Gene Synthesis services (Cat#SC1010).

### T-DNA construct and bacteria preparation

The DNA fragment of the synthetic sgRNA expression cassettes was PCR amplified with high fidelity Taq DNA polymerase (NEB Phusion^®^ Cat#M0530S) using the forward and reverse primers 5’-aggctcccgggtgcgtcgacggtctcaggtcagagcttg-3’ and 5’-gaaagctgggtgattcaagcttggtctcatcagggatccaaaag-3’ respectively. The PCR amplified DNA fragment was assembled in In-Fusion reaction (Takara^®^ In-Fusion^®^ HD Eco-Dry™ Cloning Plus Cat#638915) with a pDONR vector linearized with SalI (NEB, SalI-HF^®^ Cat#R3138S) and HindIII (NEB, HindIII-HF^®^ Cat#R3104S) to form the donor vector (pDONR-CE). The pDONR-CE vector contains the Gateway (GW) cloning sites AttL1 and AttL2. The sgRNA expression cassettes stacking was inserted between the two GW cloning sites after the In-Fusion assembling. Further through GW LR cloning reaction (Thermo Fisher Scientific ^®^, Gateway™ LR Clonase™ Enzyme mix Cat#11791019), the entire sgRNA expression cassettes fragment was transferred into a binary vector (pCUT3) which also encodes Cas9 enzyme conjugated with nuclear localization signal (NLS) and an epitope tag (HA) under the control of a UBQ10 promoter.

### Plant transformation, selection and handling

All plants used in this study were Columbia-0 (Col-0) background. All transgenic plants were transformed with the pCUT3-CE binary vector via standard Agrobacterium-mediated floral dipping as previously described. To select transgenic seeds, T1 seeds were sown on sterile Petri dishes containing Murashige and Skoog (MS) medium 0.7% (w/v) phyto-agar (Plant Media Cat#40100072-2) and 50 µg/mL Kanamycin (Fisher Scientific® CAS#25389-94-0). Seeds were surface sterilized with 10% (v/v) bleach and 0.1% Tween 20 (v/v). After 14 days of growth on sterile agar plate, resistant seedlings were scored and transferred to individual pots containing soil and were grown at 24C/16h light/100uM/cm2/sec^−1^ for further analysis and seed propagation. Green leaf tissue was collected 4 weeks later from each independent transgenic plant and stored at −80C for western blot analysis. T2 seeds were collected from individual T1 plants for further analysis. To visually select the glabrous plants, T2 seeds were stratified in 0.1% phyto-agar at 4°C for 3 days. Then, each individual seed was sown on commercial soil (50% Fafard 50% MetroMIX 360, SunGro® Horticulture) 1 cm apart from each other to facilitate visual inspection. After 3 weeks and 16hr light photoperiod at 24C, glabrous plants were visually identified by the lack of leaf trichomes. The visual identification of *jar1* and *ein2* homozygous mutants was carried out in sterile petri dishes with MS medium supplemented with ACC (Millipore Sigma^®^, SKU#A3903) or Methyl-Jasmonate (Millipore Sigma^®^ SKU#W341002) in tissue culture plates as previous described (Alonso *et al.* 1999; Staswick *et al.* 2002). Visually identified T2 seedlings were transferred to individual soil pots for further analysis and seeds propagation.

### Protein and Western Blot assay

Cas9 expression was assessed via Western blot in leaf samples. Total protein samples were extracted from 4-weeks old green leaf tissue via grinding in liquid nitrogen with mortar and pestle and resuspended in protein loading buffer (Tris-HCl, pH:8.8) and heat to 100°C for 5min. Extracts were centrifuged at 17,000x g for 5min at 25°C and the supernatant was used for gel blot analysis. Protein were subject to electrophoresis in poly-acrylamide (0.375M Tris-HCl pH=8.8, 8% Acrylamide, 0.05% APS, 0.1% SDS) matrix and transferred to PVDF membranes (Thermo Scientific™, Cat#88520) with 70V under 4°C for 90min. After blocking in milk solution, the membranes were incubated with monoclonal rabbit anti-HA antibody (Cell Signaling Technology™, mAb#3724, 1:4000 dilution) and monoclonal mouse anti-β-Actin antibody (Millipore Sigma™, Cat#MAB1501). Secondary HRP conjugated anti-rabbit IgG antibody (Jackson ImmunoResearch Laboratories, Inc. Code#111-0350144) and fluorescent conjugated anti-mouse antibody (LiCor™ IRDye^®^ 800CW Goat anti-rabbit IgG P/N#925-32210) were used for probing the primary antibody and blot visualization. ECL (Bio-Rad®, Cat#1705060) chemiluminescence and fluorescence were detected with Bio-Rad® ChemiDoc MP^®^ system and image result was analyzed by Bio-Rad Image Lab™ software.

### Gene sequencing and allele detection

To detect CRISPR-Cas9 introduced mutation, gene specific primers annealing at ∼300bp upstream and ∼300bp downstream the target site were used for PCR amplification using genomic DNA from visually selected plants and Phusion® High-Fidelity DNA Polymerase (New England BioLab^®^ Inc., Phusion®, Cat#M0530S) reaction. After the PCR, excess non-incorporated primers were removed with a single strand DNAase exonuclease (Fisher Scientific^®^ ExoSap-IT™ Cat#78-201-1ML), individual forward and reverse primers were added to separate Sanger sequencing reactions. Sanger sequencing was performed at Eurofin Genomic (Eurofins, Louisville, KY). The sequencing results were analyzed using BLAST web tool at NCBI website, aligned against wild-type (Col-0) genomic sequence. Fluorescence chromatograms were analyzed using the online tool ICE (Synthego^®^, ICE® Analysis; https://ice.synthego.com) to infer allelic composition of each T2 plant as described previously (Hsiau *et al.* 2019).

### Analysis of sgRNA nucleotide composition and secondary structure

The calculation of G/C content of all sgRNAs that were used in the experiment were done by a Python script. The secondary structure was predicted by input the sgRNA FASTA sequence into the Mfold web server (http://unafold.rna.albany.edu) as previously described (Zuker 2003).

### Data and materials availability

*E. coli* and *Agrobacterium tumefaciens* strains harboring CRISPR-Cas9 plasmids, as well as transgenic Arabidopsis plants are available upon request. The authors assure that all data necessary for confirming the conclusions are present within the article, figures, and tables. Supplementary figures available at figshare: TBD

Figure S1 shows the DNA sequence containing the three sgRNA and regulatory sequences to target *JAR1, GL1* and *EIN2*. Figure S2 shows the general scheme used for the transformation of T0 plants, the selection of T1, and the identification of T2 mutants. Figure S3 shows western blot and quantification of Cas9 in four independent T1 plants. Figure S4 shows the screening of T2 plants based of the proxy phenotypes. Figure S5 shows the secondary structures predicted for each of the three sgRNAs used in this study.

## Results

### Design of an effective CRISPR/Cas9 co-editing proxy vector

A proxy-based CRISPR-Cas9 co-editing vector is composed of: 1) Cas9 nuclease coding sequence, 2) validated sgRNA against a proxy gene driven by the RNA polymerase-II promoter, 3) sgRNAs against all genes of interest, and 4) plasmid backbone encoding all elements required for Agrobacterium mediated transformation and selection of transformants (REF). To assemble each component of the multiplex gene targeting CRISPR-Cas9 system we used the all-in-one vector pCUT3-CE (Figure 1A)(Peterson *et al.* 2016). The resulting multiplex editing vector contains: 1) Cas9-HA fusion downstream of the UBQ10 promoter and upstream of NOS terminator; 2) three sgRNA expression cassettes within which each sgRNA is flanked by a U6 promoter (AtU6) and a transcriptional termination signal (“TTTT”) to provide similar expression levels across all three sgRNA. In addition, the RNA polymerase-II transcription start site “G” was added 23 nucleotides downstream of the AtU6 promoter TATA box to efficiently initiate the transcription of each sgRNA (Figure S1); 3) all regulatory elements in the pCUT3 binary vector that allow for *E. coli* and Agrobacterium replication and provide Kanamycin resistance for the selection of transgenic T1 plants. To enable visual identification of CRISPR-mutagenized T2 plants we chose to mutagenize *GL1* (At3g27920), *JAR1* (At2g46370), and *EIN2* (At5g03280). Each of these loci reside in a different chromosome, their functions are unrelated to each other, and their loss-of-function mutants can be visually identified. The 20nt crRNA sequences were designed to target either the 1^st^ or 2^nd^ exon of each gene to increase the likelihood of yielding a null allele as a consequence of NHEJ repair generating a frame-shift or a premature stop codon for all potential isoforms of each gene (Figure 1B-D).

**Figure 1.**
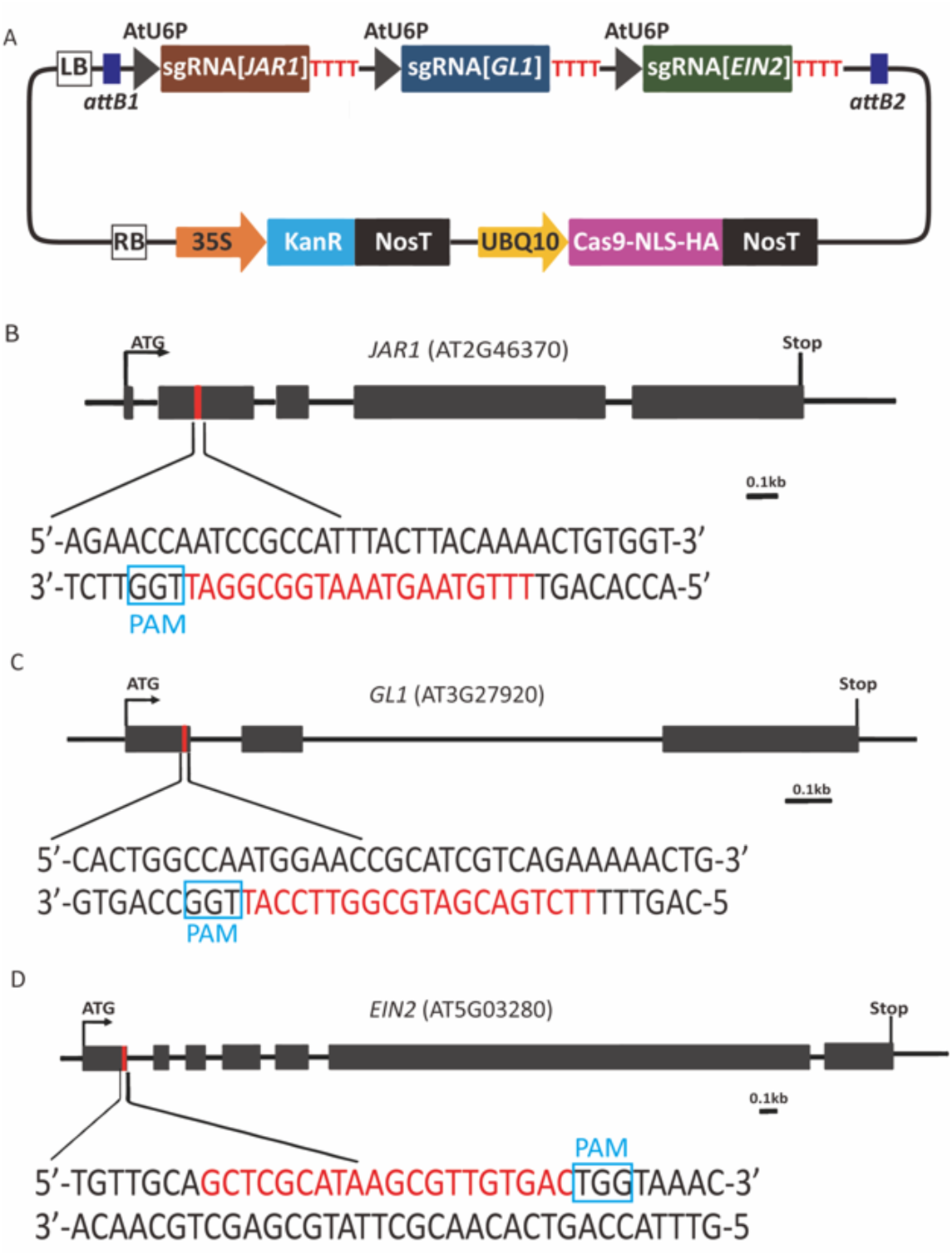
Proxy co-editing CRISPR/Cas9 construct. **A**) The pCUT3-CE construct contains three individual transcriptional units that generate sgRNA for *JAR1, GL1* and *EIN2* editing. The expression of each sgRNA is controlled by individual AtU6 promoters (U6P) and poly-T terminator (TTTT). **B-D**) *JAR1, GL1* and *EIN2* target sites. Sequence in red denote the 20nt crRNA target site within each proxy. The PAM site is boxed in blue. Scale bar=0.1kb

### Heritable CRISPR-Cas9-mutageniced proxies are easily detectable in T2 progenies

After Agrobacterium-mediated transformation of immature flowers of Col-0 wild type plants, we selected for transformants on agarose plates using Kanamycin 50 µg/mL as the selective agent. From five independent transformation experiments, we recovered a total of twenty-six T1 transgenic plants. After self-fertilization, we randomly chose the progeny of four T1 lines (#1, #3, #4 and # 25), obtained from four independent transformation experiments, for proxy-based visual screening of CRISPR-edited T2 plants (Figure S2). For each of the four independent T1 lines, we grew ∼1000 T2 seeds to screen for loss of function of proxies (*jar1, gl1 and ein2 plants*). Among T2 plants visually inspected for lack of trichomes in the progeny of T1 lines #1, #3 and #25, *gl1* mutants appeared with frequencies ranging from ∼2 to 6.3 % (Figure 2A), suggesting a high-efficiency of editing. For *jar1* or *ein2* selection, we sown ∼1,000 T2 seeds on Murashige and Skoog phyto-agar plates containing either the JA-precursor Methyl Jasmonate (MeJA) or the ETprecursor Amino-Cyclopropane-Carboxylic acid (ACC), respectively. One week after germination on MeJA-containing square plates incubated vertically to allow for root length assessment, wild type seedlings produced roots of 0.5±0.2 cm in length, while *jar1* positive control seedlings produced roots of more than 1.5 cm in length. T2 seedlings from individual T1 progenies with roots of more than 2 cm in length were selected for further analysis. Five days after sowing on ACC-containing plates incubated in the dark, wild type seedlings produced hypocotyls of approximately 0.5 cm in length with the typical downward cubature, while *ein2* positive control seedlings produced straight hypocotyls of more than 1 cm in length. T2 seedlings from individual T1 progenies with straight long hypocotyls were selected for further analysis. Again, we found *jar1* and *ein2* plants in the progeny of T1 lines #1, #3 and #25, but we did not identify mutants in the T2 progeny of T1 line #4 (Figure 2). Western blots of leaf protein samples of each T1 line analyzed showed Cas9 expression in lines #1, #3 and #25, but no in T1 line #4, explaining the lack of observable phenotypes in the T2 progeny of this T1 line (Figure 3). As expected, each individual T1 plant expressed different levels of Cas9, a variation that may stem from the independent chromosomal location of the T-DNA in each T1 plant analyzed. Interestingly, the percentage of visually identified T2 plants varied across independent T1 lines but did not correlate with Cas9 expression. The frequency of *jar1* and *ein2* plants in T2 progenies ranged from 0.001 to 0.002% and 0.9 to 1.8% for *jar1* and *ein2*, respectively (Figure 2B and 2C). The lower recovery of *jar1* and *eni2* relative to *gl1* mutants may in part relate to the quantitative scoring of these phenotypes (as opposed to qualitative scoring of *gl1*), and to our stringent cutoff for root and hypocotyl length (see above).

**Figure 2.**
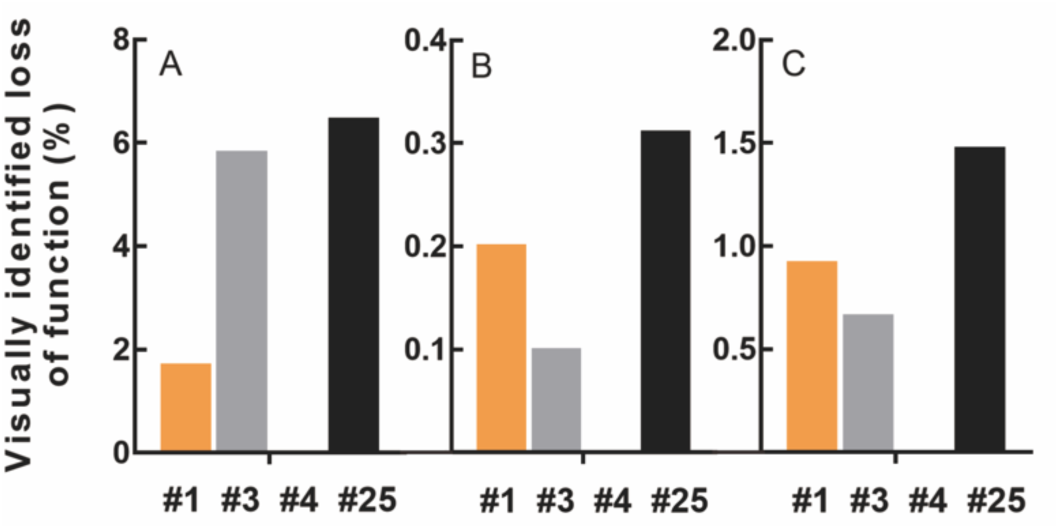
Frequency of loss-of-function mutants visually identified in T2 progenies. Percentage of *gl1* (**A**), *jar1* (**B**) and *ein2* (**C**) plants across T2 progenies of three independent T1 lines (#1, #3 and #25) calculated as number of plants with mutant phenotypes/total number of T2 plants obtained from each independent T1 plants (#1, #3, #4 and #25). Total number of *gl1, jar1* and *ein2* recovered was 130, 5 and 32 plants, respectively.

**Figure 3.**
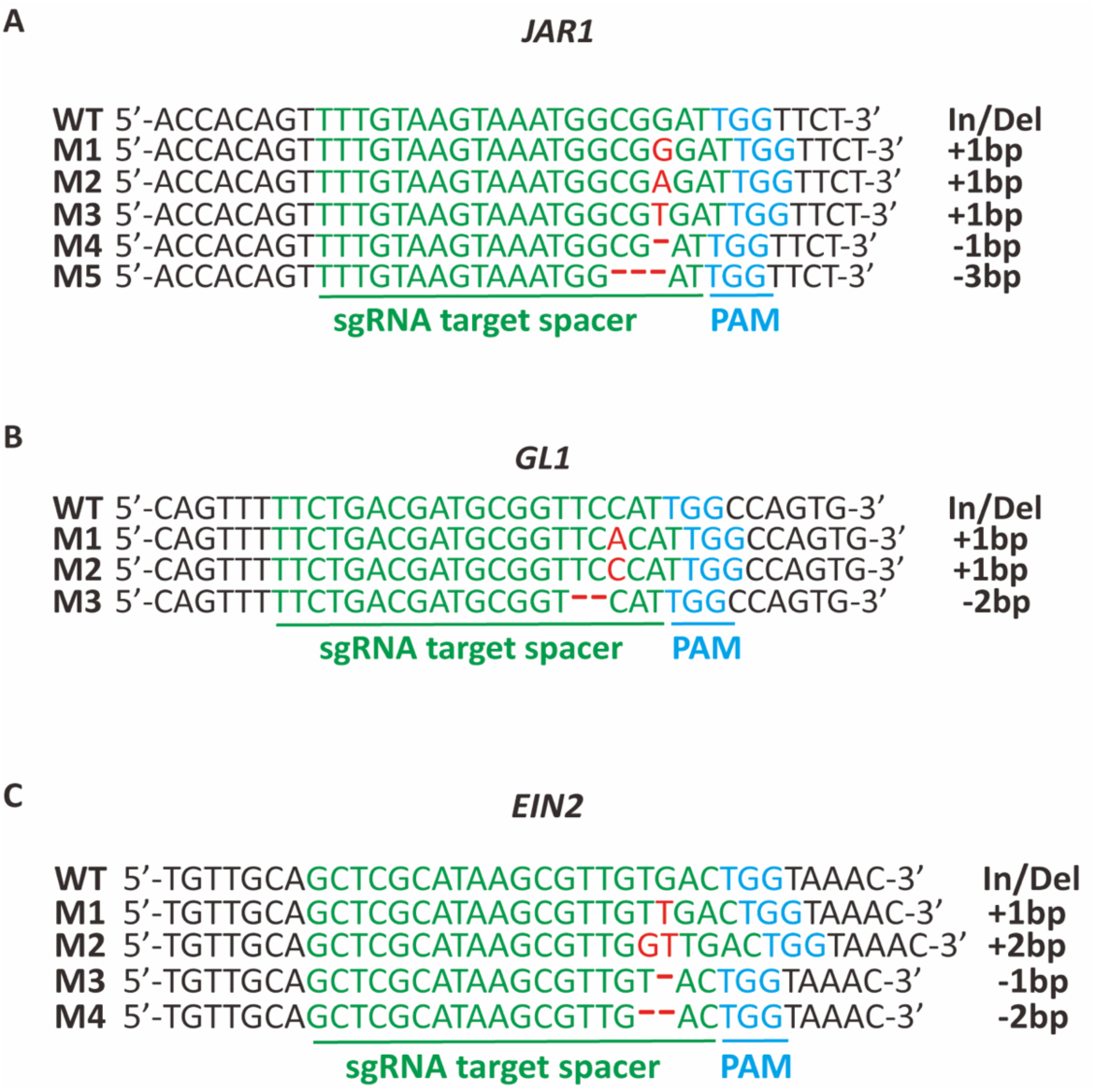
Sequence analysis of CRISPR/Cas9 edited proxy genes. A-C) CRISPR/Cas9 induced mutations detected via PCR and Sanger sequencing in *jar1* (**A**), *gl1* (**B**) and *ein2* (**C**) mutants across independent T2 progenies. The target sequence is depicted in green. The PAM site is depicted in blue. Indels are depicted in green. Wild type reference DNA sequence (WT) and mutant alleles (M1…5) detected for each proxy gene are shown as sequence alignments.

Due to the UBQ10 promoter activity, the constitutive and ubiquitous expression of Cas9 may produce edited cells or lineages in somatic and/or germline tissues, yielding mosaic T2 plants. To reduce the odds of analyzing mosaic edited plants, we only selected T2 plants that had no trichomes anywhere in the leaves, or showed long roots (more than 1.5 cm) or long straight hypocotyl (Figure S4). We reasoned that mutations in these T2 plants should have originated in T1 germline and hence their genetic makeup should be more uniform across reproductive and somatic tissues. To assess the genetic makeup of the selected mutants, plants were allowed to self-pollinate and produce T3 seeds for progeny tests. All the T2 plants that show *jar1, gl1* or *ein2* phenotypes produced 100% *jar1, gl1* and *ein2* mutant plants in their T3 progenies, respectively, which confirmed that the selected T2 plants were loss-of-function mutants, as indicated by their phenotypes.

### Sub-optimal sgRNA secondary structure may limit the efficiency of CRISPR-Cas9 editing

To understand the underlying causes that produced different mutagenesis frequencies of the three targeted loci, we investigated three important factors related to sgRNAs design that are known to impact CRISPR editing: 1) number of crRNA off targets; 2) crRNA GC content; 3) sgRNA secondary structure. The first factor under consideration, the specificity score, which is reported as a whole number out of 100 perfect score, the values of the three 20nt crRNA sequences were similar (cr*JAR1*: 98/100, cr*GL1*: 98/100, cr*EIN2*: 99/100). Yet, the number of potential off-targets within the Arabidopsis genome were higher for *JAR1* crRNA than for *GL1* and *EIN2* crRNAs (cr*JAR1*: 9, cr*GL1*: 5, cr*EIN2*: 3). Assuming that there is competition for sgRNAs across potential off-targets at different loci, we would expect *EIN2* or *JAR1*, the outermost sgRNA in the array, to show the highest efficiency. In addition, the numbers of off-targets do not seem to explain the high mutagenesis of *GL1* and the low mutagenesis of *JAR1* (Doench et al., 2016). Therefore, we should consider other explanations. The G/C content of the three crRNAs varied substantially, being 35% for cr*JAR1*, 50% for cr*GL1*, and 55% for cr*EIN2*, suggesting that the low G/C content of cr*JAR1* could, at least in part, explain its low mutagenesis efficiency. However, the GC content for all 3 crRNAs falls within the effective range of 30 to 80% (Liang *et al.* 2016), and the GC content of cr*GL1* and cr*EIN2* are only 5% different while the editing efficiency of sg*GL1* is up to 6-fold higher than that of sg*EIN2*. Therefore, GC content does not seem to be a major determinant of the efficiency differences across the three sgRNAs. A third factor to consider is the secondary structure of the sgRNA. Previous studies revealed that three stem loops (hairpins) are necessary for the effectively formation of a DNA-sgRNA-Cas9 complex (Liang et al., 2016). Among these, stem loop #1 is crucial for the formation of a functional Cas9-sgRNA-DNA complex, while stem loop #2 is critical to improve complex stability and *in vivo* activity. We performed *in-silico* analysis of the *JAR1, GL1* and *EIN2* sgRNAs sequences using the online tool Mfold (Zuker et al., 2003). A single *in-silico* prediction revealed that all three stem loops are intact in the *GL1* sgRNA (Figure S5D). The *EIN2* sgRNA analysis rendered two predictions (Figure S5E-F), one showing all three intact stem loops and one showing a missing stem loop #1. The secondary structure predictions of *JAR1* sgRNA showed that stem loops #1 and #2 are missing in all three predicted structure (Figure S5A-B-C). Hence, the presence of three stem loops and the low number of CBPs and IBPs in *GL1* sgRNA may explain the high efficiency of *GL1* mutagenesis. The presence of two stable and one unstable stem loop, together with the higher number of CBPs and IBPs, may explain the intermediate efficiency of the *EIN2* sgRNA. Finally, the absence of stem loops #1 and #2, and high number of CBPs and IBPs would explain the low efficiency of *JAR1* sgRNA. Therefore, an optimal sgRNA secondary structure seems critical for effective CRISPR-Cas9 editing, and may be carefully considered at the time of designing sgRNAs.

### Efficient identification of Cas-9-editing at other genomic loci using proxy-based selection

For most gene-editing endeavors, the phenotype/s associated to the gene/s of interest is not known. In fact, the purpose of using CRISPR-editing in most cases is to uncover phenotypes associated to the function of the gene/s of interest. To verify the mutations at each locus, we analyzed the DNA sequences of visually selected *gl1, jar1*, and *eni2* plants. We collected leaf samples of *gl1, eni2* and *jar1*-selected T2 plants, extracted DNA, and PCR-amplified and Sanger-sequenced 600bp of DNA surrounding the crRNA target sequence of the three proxy genes. A CRISPR-edited locus typically consists of a mix of mutant and wild type sequences (alleles). This complexity stems from DSB of DNA introduced by Cas9 followed by DNA repair via NHEJ, a process that may start when the T-DNA harboring CRISPR-Cas9 is integrated into the genome of germline cells in T0 plants (see Figure S1). Throughout the T1 and T2 generations, Cas9 will continue to edited wild type target sequences in germline as well as somatic cells. The analysis of the DNA sequences was accomplished using the ICE (Inference of <u>C</u>RISPR <u>E</u>dits) sequence analysis tool (https://ice.synthego.com). ICE weighs the quality of each DNA sequence and determine the relative representation of each of the four possible nucleotides at each given position in the DNA sequence via assessing the area below the fluorescence chromatogram peak at each position (Hsiau *et al.* 2018). Among the visually selected *gl1, jar1* and *eni2* plants, we identified three different alleles of *gl1*, five alleles of *jar1*, and four alleles of *eni2* (Figure 3). The mutations detected in the three proxies were consistent with previous studies reporting insertions of 1 or 2 nucleotides (+1 or +2) and deletions of 1, 2 or 3 nucleotides (−1, −2 and −3) within the six nucleotides upstream of the PAM sites, which are the hallmarks of NHEJ-mediated DNA repair in Arabidopsis (Jinek *et al.* 2012; Oost 2013; Peterson *et al.* 2016).

To assess the power of each of these proxies to predict coediting, we used PCR and Sanger DNA sequencing analyses to detect mutant alleles in surrogate genes; for *gl1* selected T2 plants we PCR and sequenced *JAR1* and *EIN2* targets, for *ein2* selected T2 plants we PCR and sequenced *JAR1* and *GL1* DNA targets, and for *jar1* selected T2 plants we PCR and sequenced *EIN2* and *GL1* targets. We scored the frequency of single and double co-editing, indicative of plants carrying mutant alleles in two or three of the targeted genes, respectively, via counting the number of plants with unresolved DNA sequences upstream of the PAM site of each target divided by the total number of T2 plants visually identified with each proxy (smooth leaves, JA insensitivity, or ET insensitivity). The frequency at which each mutant allele was detected varied across the T2 progenies of independent T1 plants. Among plants without trichomes (*gl1* = smooth leaves), the frequency of single co-editing varied from 61.1% to 93.75% for either *JAR1* or *EIN2* (Figure 4). Further, the frequency of double co-editing (yielding a triple *gl1*-*jar1*-*eni2* mutant) ranged from 18.5% to 50% when using *gl1* as the proxy (Figure 4A). Among ET insensitive plants (*eni2*), single gene co-editing frequency varied from 55.6% to 81.25%, while double co-editing ranged from 0% to 56.25% (Figure 4B). Although JA insensitive plants (*jar1*) were recovered through *in plate* selection of T2 progenies from the three T1 plants expressing Cas9, we only recovered a total of five *jar1* plants (2 from T1 #1, 1 from T1 #3, and 2 from T1 #25). Therefore, the frequency of co-editing cannot be accurately estimated and was not included in Figure 4. Nevertheless, every one of the five *jar1* T2 mutants recovered were also mutants for both *GL1* and *EIN2*, suggesting high efficiency of co-editing (up to 100%) when using jar1 as the initial selection proxy.

**Figure 4.**
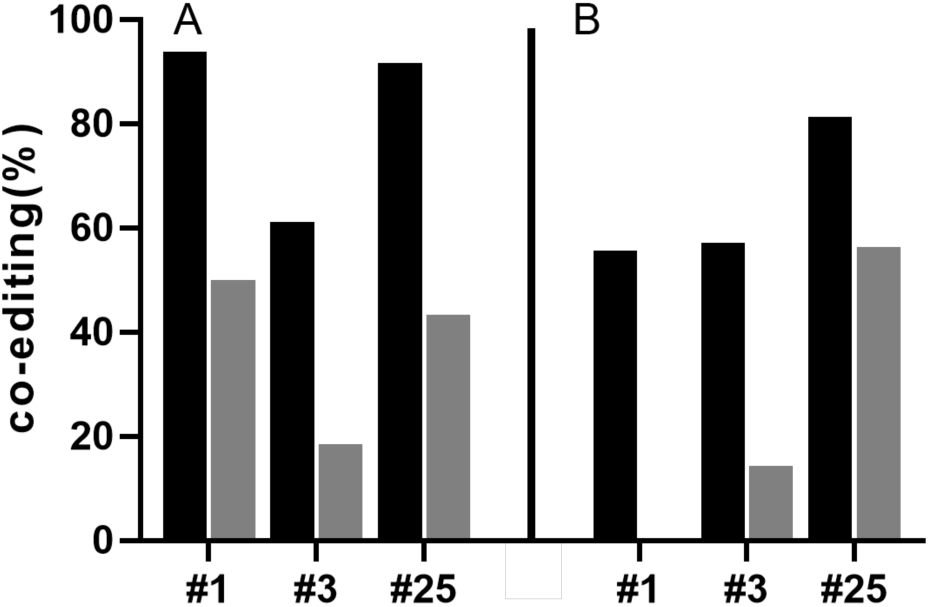
Predictive power of proxy genes. Scoring of mutant alleles detected in one (black) or two (grey) surrogate gene in addition to the loss-of-function proxy. Visual selection of proxy based on **A**) lack of trichomes (*gl1*) or **B**) ethylene insensitivity (*ein2*).

A detailed analysis of the DNA sequence of each surrogate gene using the ICE online tool revealed a complex allelic composition in the somatic cells of most T2 plants. As expected, we detected wild type, homozygote mutants, heterozygotes and Bi-allelic (or higher order allelism) sequences for each gene under study across different plants (Feng *et al.* 2014). Thirty percent of co-edited plants initially selected using *gl1* did not have wild type alleles of *JAR1* or *EIN2*. Triple mutants were found in the T2 progeny of T1 line #25, where wild type alleles for *JAR1* and *EIN2* were not detected in 3.85% of plants analyzed. Importantly, up to 30.8% of the T2 plants in this progeny did not have wild type alleles for a second gene and had a mix of wild type and mutant alleles for a third targeted gene (Table I). Among *ein2* plants, we found up to 46.2 % of plants (in T2 progeny of T1 #25) that were also homozygous for either *jar1* or *gl1* mutant alleles. Most *ein2* mutants in every of the three T2 progenies analyzed were heterozygous for either *JAR1* or *GL1* (Table I). Interestingly, neither the percentage of visually selected plants (Figure 2) nor the percentage of plants with mutant alleles in a second or a third gene within the visually selected T2 plants (Figure 4) seem to correlate with Cas9 expression (Figure S3).

**Table I.**
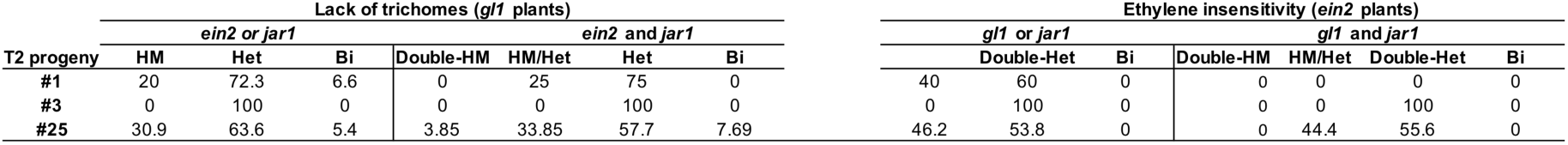
Allelic constituency of co-edited genes. Numbers indicate the percentage of co-edited surrogate genes in T2 plants of three independent progenies (#1, #3 and #25) and their allelic state. HM: homozygous. HET: heterozygous. Bi: bi-allelic. Double-HM: no wild type alleles for both genes under study. HM/HET: no wild type alleles for one gene and one wild type allele for the second gene under study. Double-HET: one wild type allele for each gene under study. Bi-HET: multiple alleles in one gene and at least one wild type allele for the second gene under study.

Overall, our data demonstrate that proxies of CRISPR mutagenesis are powerful predictors of the occurrence of CRISPR-induced mutations in surrogate genes, thus allowing for the rapid identification of plants where genes of interest have been edited as well.

## Discussion

The challenge associated to the identification of a handful of edited plants among hundreds of non-edited plants was recognized early when CRISPR-Cas9 genome-editing was proven to work in plants. Previous studies have used fluorescent protein-Cas9 fusions or epitope-tagged Cas9 to identify T1 plants that express Cas9, which has significantly reduced the resources spent in the identification of CRISPR mutagenized plants (Osakabe *et al.* 2016; Tang *et al.* 2018). Although a significant step forward, these strategies involve imaging leaf samples with a fluorescence microscope or taking leaf samples to run western blots to detect Cas9-expressing T1 plants. In addition, a large number of T2 progenies of the Cas9 expressing plants need to be DNA-profiled in order to identify the T2 plants harboring mutant alleles. Our proxy-based selection approach greatly reduces the number of T2 plants that need to be DNA profiled as 40-100% of proxy-based selected plants had mutations in a second or a third gene of interest. As for the most effective manner to apply proxy-based selection, some considerations may be necessary. For instance, the most effective sgRNA was the one targeting *GL1*. However, when looking at the frequency of co-editing, although high (∼80% of the *gl1* plants had mutations in a second gene, and ∼40% had mutations in a second and a third gene of interest), *gl1* was a less effective proxy than *jar1*, as all five *jar1* plants isolated in this study had mutations in both *GL1* and *EIN2*. It is tempting to speculate that the high frequency of co-editing observed in *jar1* mutants could be due to the low mutagenesis observed in *JAR1* combined with higher mutagenesis in *GL1* and *EIN2*. Our analysis showed that the mutagenesis efficiency of each gene tested correlates with its sgRNA secondary structure, which in turns is highly dependent on the gene-specific crRNA sequence. In light of this observation, it seems reasonable to hypothesize that using a suboptimal (low efficiency) sgRNA to target the proxy of choice, in combination with highly efficient sgRNA/s to target the gene/s of interest, could dramatically reduce the number of plants that need to be DNA profiled to identify CRISPR multiplex mutants. Our results, in combination with previous reports showing that up to several genes can be Cas9 mutagenized simultaneously, support the notion that our approach could be scaled up to target more than three genes at a time (Feng *et al.* 2014; Zhang *et al.* 2016).

Importantly, once the T2 plants of interest are identified, progeny tests or genotyping of T3 progenies allows for the selection of mutant plants that no longer harbor the T-DNA. This would prevent the continuous mutagenesis of the target genes by the ubiquitously expressed Cas9, and minimize the likelihood of off-target mutagenesis. In cases where the proxy represents an impediment for the analysis of gene function, the mutants could be crossed with wild type plants to segregate the proxy and the T-DNA from the mutations of interest and select for F2 plants that express a wild type version of the proxy and no longer express CRISPR-Cas9. *JAR1, GL1* and *EIN2,* are encoded in chromosomes 2, 3 and 5, respectively, enabling to choose a proxy gene that is not linked to the gene/s of interest, and can later be easily segregated out. While outcrossing the mutants, it is important to select for plants that no longer express Cas9.

Unlike Cas9 fusions to fluorescence proteins or epitope tags, the identification of *jar1, gl1* and *ein2* mutant plants does not require any sophisticated piece of equipment. While *jar1* and *ein2* plants can be identified in culture plates within 1 week after germination, the identification of g*l1* mutants only involves growing T2 plants in soil for three weeks and visually identifying plants with smooth leaves among hairy wild type plants, thus, making this strategy more accessible for laboratories with little resources or even the classroom setting. The strategy laid out in this study constitutes a proof-of-principle of the potential use of proxies to facilitate the selection of genome-edited plants. The choice of the appropriate proxy will depend on the plant species and on the phenotypes expected/predicted to be associated to the gene/s of interest. In any event, *GL1* mutant alleles are found in many naturally occurring ecotypes of Arabidopsis, which suggests that mutation in *GL1* should not produce overly pleotropic phenotypes that could interfere with the function of the genes of interest that are targeted simultaneously, unless of course, they also affect trichome development. Similar selection strategies could be used for the identification of CRISPR-Cas9 mutagenized plants in crops with more complex, often polyploid genomes, where the identification of multiplex mutants remains challenging.

## Acknowledgments

We express our gratitude to Michael P. Timko and Keith G. Kozminski for their advice during the execution of this study, and to David M. Parichy for reading the manuscript.

