## Supplementary Figure S1-S5 for "Proxies of CRISPR-Cas9 activity to aid in the identification of mutagenized Arabidopsis plants"

GGTCTCAGGTCAGAGCTTGTTCAGGACTCGAGcatcttcattcttaagatatg
aagataatcttcaaaaggcccctgggaatctgaaagaagagaagcaggcccatttatatgggaaagaaca
atagtatttcttatataggcccatttaagttgaaaacaatcttcaaaagtcccacatcgcttagataagaaaacg
aagctgagtttatatacagctagagtcgaagtagtgattg**TTTGTAAGTAAATGGCGGA**
**T**GTTTTAGAGCTAGAAATAGCAAGTTAAAATAAGGCTAGTCCGTT
ATCAACTTGAAAAAGTGGCACCGAGTCGGTGCTTTTGAGCTCcatc
ttcattcttaagatatgaagataatcttcaaaaggcccctgggaatctgaaagaagagaagcaggcccattta
tatgggaaagaacaatagtatttcttatataggcccatttaagttgaaaacaatcttcaaaagtcccacatcgc
ttagataagaaaacgaagctgagtttatatacagctagagtcgaagtagtgattg**TTCTGACGAT**
**GCGGTTCCAT**GTTTTAGAGCTAGAAATAGCAAGTTAAAATAAGG
CTAGTCCGTTATCAACTTGAAAAAGTGGCACCGAGTCGGTGCTTT
TGAGCTCcatcttcattcttaagatatgaagataatcttcaaaaggcccctgggaatctgaaagaagag
aagcaggcccatttatatgggaaagaacaatagtatttcttatataggcccatttaagttgaaaacaatcttcaa
aagtcccacatcgcttagataagaaaacgaagctgagtttatatacagctagagtcgaagtagtgattg**GC**
**TCGCATAAGCGTTGTGAC**GTTTTAGAGCTAGAAATAGCAAGTTA
AAATAAGGCTAGTCCGTTATCAACTTGAAAAAGTGGCACCGAGT
CGGTGCTTTTGGATCCCTGATGAGACC

**Figure S1**. In-silico designed and *in vitro* synthesized sgRNA expression cassettes stacks. Each AtU6 promoter sequence is depicted in lowercase. Each sgRNAs is depicted in uppercase brown (*JAR1*), blue (*GL1*) and green (*EIN2*). Each crRNA sequence is depicted in bold font. The RNA polymerase III transcription start site “g” is depicted in lower case red font and the stop site (poly T tail) in uppercase red font. The 32 and 17 extra nucleotides at 5’ and 3’ of the synthetic DNA fragment are shown in black uppercase font.

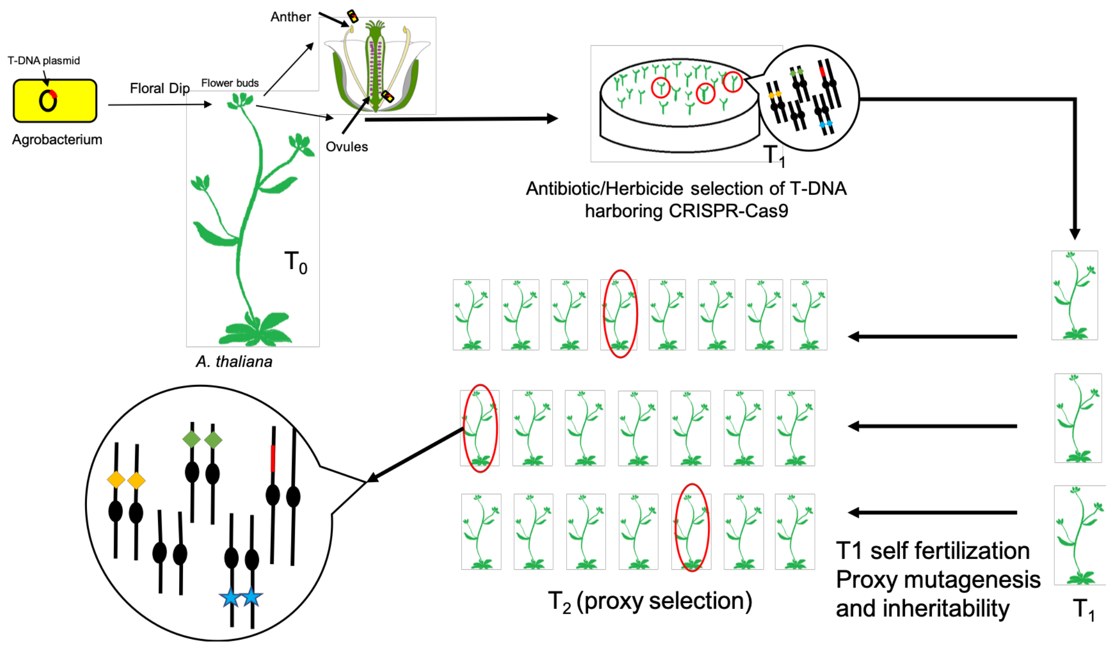


**Figure S2**. Selection scheme of CRISPR-mutagenized plants. In the T1 generation, each gene targeted is depicted as a yellow, green or blue diamond each one positioned on different Arabidopsis chromosomes. The T-DNA harboring CRISPR-Cas9 is depicted as a red chromosome fragment. The genotype of proxy-selected T2 plants is shown at the end of the scheme: the blue star depicts loss-of-function mutation of the proxy gene, while yellow and green diamonds depict surrogate genes of unknow condition.

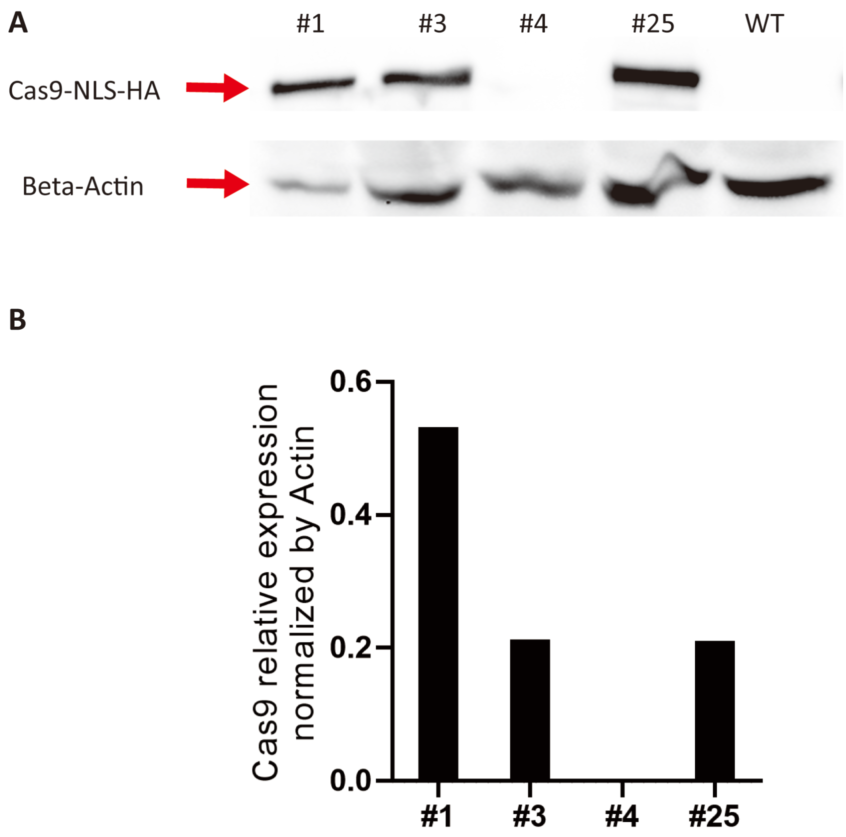


**Figure S3**. Cas9 expression. A) Cas9 expression in 4 independent transgenic lines (#1#3#4#25) and non-transgenic wild type plants (WT) were tested via western blot. B) Quantification of Cas9 expression normalize by Beta-Actin. This experiment was repeated three times with similar results.

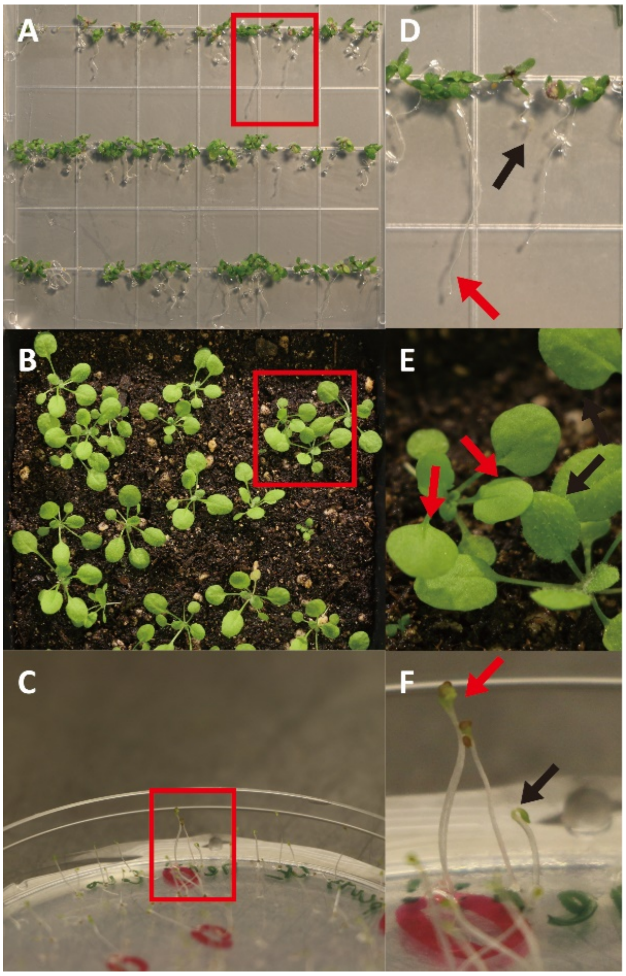


**Figure S4**. Visual identification of CRISPR-Cas9 mutants in T2 progenies. **A-C)** Proxy phenotypes for *jar1* (**A**), *gl1* (**B**) and *ein2* (**C**) in T2 progenies. The red inset highlights proxy plants side-by-side with plants showing wild type phenotype. **D-F**) Closeups of *jar1* (**A**), *gl1* (B) and *ein2* (**C**) proxy plants. Red and black arrows in D-E-F indicate mutant and wild type phenotypes for each proxy.

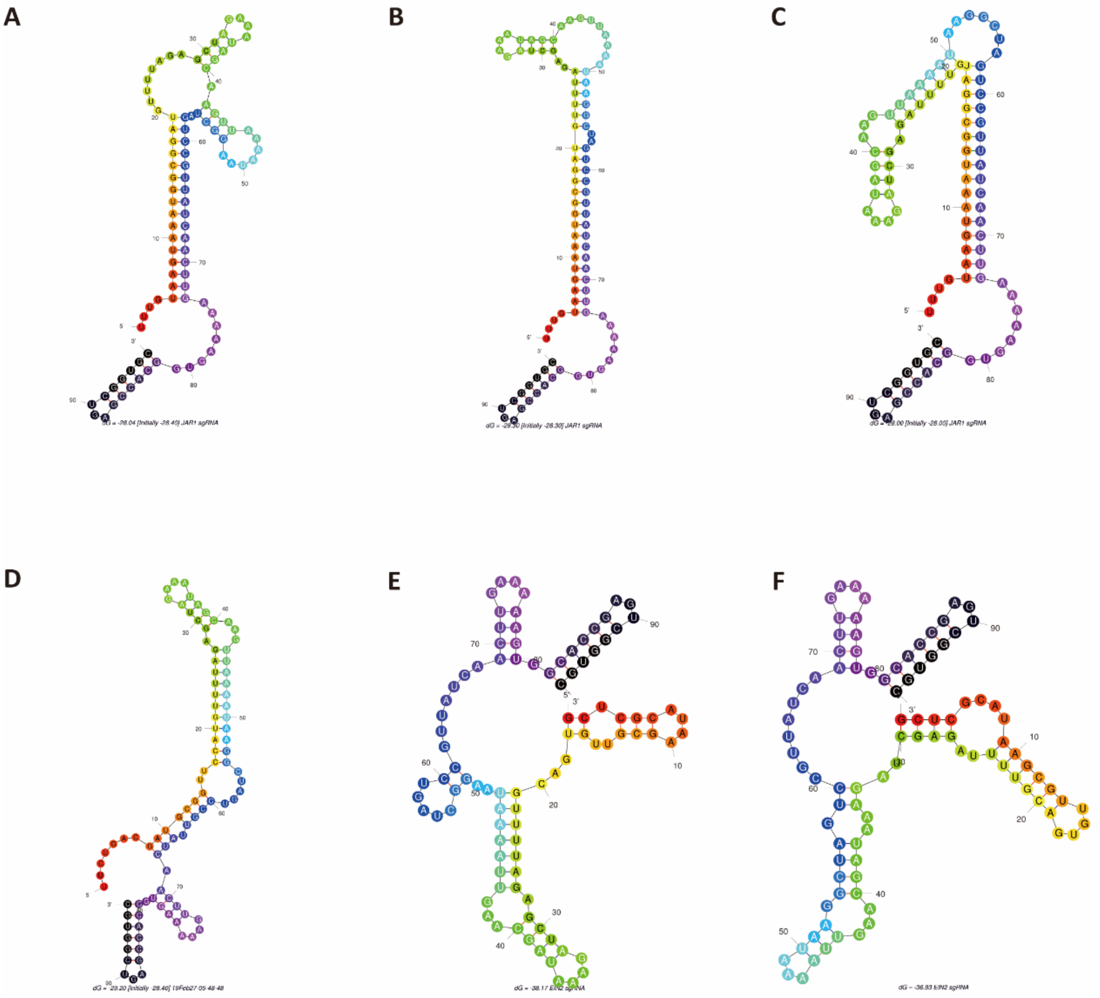


**Figure S5**. sgRNA secondary structure. In-silicon predictions of *JAR1*(**A-C**), *GL1*(**D**) and *EIN2* (**E-F**) sgRNA secondary structure by Mfold (Zuker et al. 2003). Nucleotides were colored coded as red and black for the 5’ and 3’ ends, respectively.
